# Localizing electrophysiologic cue-reactivity within the nucleus accumbens guides deep brain stimulation for opioid use disorder

**DOI:** 10.1101/2024.12.30.630822

**Authors:** Liming Qiu, Young-Hoon Nho, Robert Seilheimer, Min Jae Kim, Altona Tufanoglu, Nolan R. Williams, Anna Wexler, David W. Oslin, Bruno Millet, Katherine W. Scangos, Bijan Pesaran, A. Eden Evins, R. Mark Richardson, Anna-Rose Childress, Casey H. Halpern

**Affiliations:** Department of Neurosurgery, Perelman School of Medicine, University of Pennsylvania, Philadelphia, PA, USA; Department of Psychiatry, Perelman School of Medicine, University of Pennsylvania, Philadelphia, PA, USA; Department of Psychiatry and Behavioral Sciences, Stanford University, Palo Alto, CA, USA; Department of Medical Ethics and Health Policy, Perelman School of Medicine, University of Pennsylvania, Philadelphia, PA, USA; Department of Psychiatry, Corporal Michael J. Crescenz Veterans Affairs Medical Center, Philadelphia, PA, USA; Department of Psychiatry, Paris Brain Institute, Pitie-Salpetrière Hospital, Sorbonne University, Paris, France; Department of Bioengineering, University of Pennsylvania, Philadelphia, PA, USA; Department of Neuroscience, University of Pennsylvania, Philadelphia, PA, USA; Center for Addiction Medicine, Massachusetts General Hospital, Harvard Medical School, Boston, MA, USA; Department of Neurosurgery, Massachusetts General Hospital, Harvard Medical School, Boston, MA, USA; Department of Surgery, Corporal Michael J. Crescenz Veterans Affairs Medical Center, Philadelphia, PA, USA

**Keywords:** Cue reactivity, Deep brain stimulation, Electrophysiological biomarker, Opioid use disorder

## Abstract

Substance use disorder (SUD) is a significant public health concern, with over 30% of the affected population not responding to available treatments. Severe SUD is characterized by drug-cue reactivity that has been reported to predict treatment-failure. We leveraged this pathophysiological feature to optimize deep brain stimulation (DBS) of the nucleus accumbens region (NAc) in an adult with SUD. A personalized drug cue-reactivity task was administered while recording NAc region electrophysiology from a lead externalized for clinical purposes. We identified a drug cue-evoked signal in the ventral NAc associated with intensification of opioid-related cravings, which attenuated subsequent to stimulation delivered to the same area. DBS was then programmed to engage this focal region, which resulted in sustained suppression of drug-related cravings. This finding heralds the potential for personalized strategies to optimize DBS for SUD.

---

Substance use disorders (SUDs) are a growing public health concern of epidemic proportion, resulting in more than 600,000 deaths in the Unites States annually, with over 100,000 annual deaths in the United States from opioid use disorders (OUD) alone^1^. The greatest challenge to treatment success is preventing relapse, and relapse risk has been attributed at least in part to neural adaptations in the frontostriatal circuitry involving the nucleus accumbens (NAc), an important component of the mesolimbic dopamine system^2^. Functional neuroimaging assays utilizing cue-reactivity paradigms to probe neural activity underlying drug desire (“craving”)^3,4^, compulsive drug seeking and drug-use behaviors have revealed hyperreactive responses to drug-related cues. These phenomena are associated with poorer treatment outcomes and increased relapse rates^5,6^. However, a well-recognized challenge with functional MRI is its limited spatial and temporal resolution, which preclude exquisite detailing of specific anatomophysiological involvement of the NAc region. Deep brain stimulation (DBS) is a long-term neuromodulatory treatment with the potential to rescue frontostriatal adaptations and reverse behavioral over-response to drug cues in severe cases^7^. However, the best way to engage the NAc network with DBS in SUD remains largely unknown^8^.

In this case study, we seized a serendipitous clinical opportunity to synchronize a personalized cue-reactive paradigm to intracranial electroencephalographic (iEEG) recordings via an *in-situ* NAc DBS electrode. This afforded high spatial and temporal resolution to decipher the involvement of the NAc region in drug-related cravings and behavior^4,9^. We report the identification and use of a specific drug cue-reactive electrophysiological biomarker to optimize DBS programming in a patient with OUD. Our work heralds the potential for personalizing DBS using an objective read-out of at-risk pathophysiology.

The patient is a 25-year-old male with an existing NAc DBS to treat his severe OUD off-label (see *Online Methods*; Fig S1a). DBS therapy improved his OUD and substance cravings, allowing him to abstain from drug use for several years, but a prohibitively high dose of stimulation (14.5mA) in the ventral anterior limb of the internal capsule (vALIC) was required. He suffered a wound erosion over the left subclavicular incision that warranted removal of the implantable pulse generator (IPG), which was followed by rapid recurrence of opioid cravings and drug-related thoughts (Fig 1a). Due to suspicion of ongoing infection, the DBS system was externalized with an extension cable while cultures and antibiotic treatment ensued (*Online Methods*).

**Figure 1.**
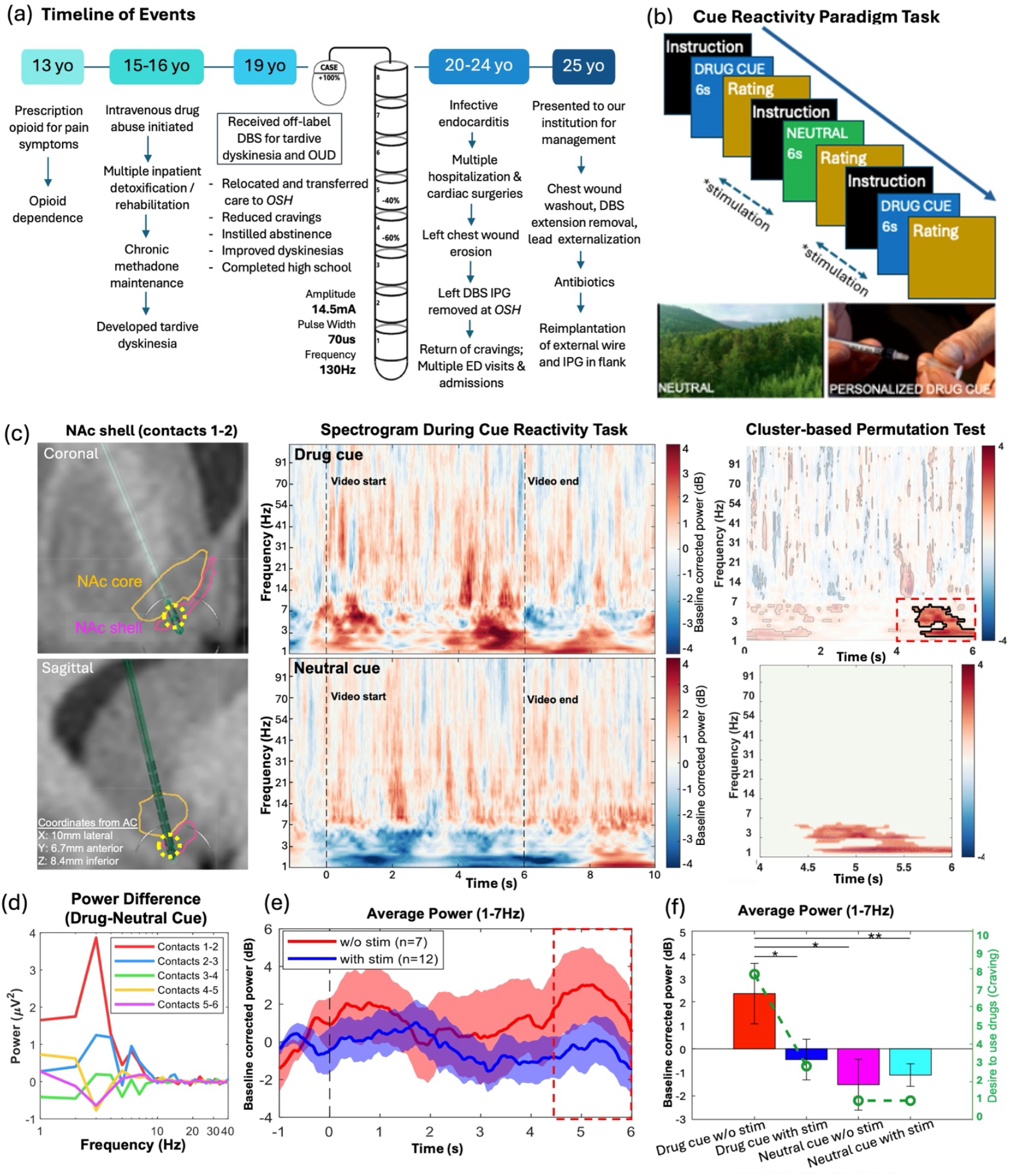
A specific 1-7Hz low-frequency electrophysiological biomarker can be identified with a personalized cue-reactivity paradigm task. (a) Timeline of events leading to subject presentation and evaluation for DBS replacement and optimization. (b) Schematic of personalized drug cue-reactivity paradigm task based on pre-task interview of the patient’s drug use habits. In stimulation trials, 10 minutes of stimulation was delivered prior to administration of the task, as well as intermittent stimulation (∼15 seconds) before each cue was presented (*). (c) A specific, low-frequency (1-7Hz) biomarker associated with drug cravings was identified in the presumed nucleus accumbens (NAc) shell subregion. Left panel; Coronal (top) and sagittal (bottom) T1 MRI views of DBS electrode contacts in the NAc shell. Yellow dotted circle demarcates contacts 1-2 where bipolar recordings were obtained. Middle panels; Time-frequency analysis (−1 to 10s; video starts at 0s and ends at 6s) showed a significant dissociation in power in the low-frequency range when drug-related cues were presented (*Top*), as compared to neutral cues (*Bottom*). Right panel; Cluster-based permutation test was used to identify time window and frequency ranges that were significantly different between drug cue and neutral cue trials (*Top*). Dotted red box (4.5 to 6s) denotes time window and cluster that survived permutation testing, showing a statistically significant increase in low frequency (1-7Hz) power upon presentation of drug-related cues (bottom). Similar analysis did not identify any significant difference in power in iEEG recorded in other electrode contact pairs. (d) Plot of power spectral density (PSD) differences in trials of drug vs neutral cues showed a large dissociation in the low frequency range, only when measured in contacts 1-2, but not in other contact pairs, suggesting anatomical specificity. (e) Average power of 1-7Hz frequency band across time, without stimulation (red) and with NAc shell stimulation (blue). Red dotted box denotes time period when power level was significantly reduced with stimulation (p=0.039). (f) Bar plot of average 1-7Hz band power measured in contacts 1-2 under four conditions. Power was significantly elevated when drug cues were presented (red; n=7) vs neutral cues (pink; n=10) without stimulation (p=0.018). With stimulation, 1-7Hz power was significantly reduced when drug cues were presented (blue; n=12) (p=0.039) and this was not statistically different from power in neutral cue condition (cyan; n=12). Average craving ratings for each condition is shown in green open circle. Stimulation of the NAc shell significantly reduced cravings levels when drug cues were presented (*p*<5×10^−7^), but did not affect ratings in neutral cues (dashed green line). *AC = anterior commissure; DBS = deep brain stimulation; ED = emergency department; IPG = implantable pulse generator; MRI=magnetic resonance imaging; NAc=nucleus accumbens; OSH = outside hospital; OUD = opioid use disorder*.

We hypothesized that prior high charge requirements were due to non-specific engagement of the NAc. To identify focal sites within the NAc region where cue-reactivity was most robust and specific to drug cues, three computer-based tasks were conducted: a personalized primary drug cue-reactivity task to assess for presence of drug-related biomarker^6^ (Fig 1b), the Monetary Incentive Delay, MID^10^ and the Blink Suppression Task^11^ (*Online Methods*, Fig S2 and S3). The latter two tasks allowed us to assess specificity of electrophysiologic drug-cue reactivity.

Presentation of personalized drug-related video cues evoked a robust elevation of a low-frequency band (1-7Hz) power in the iEEG recordings within the ventromedial-most region of the NAc, the presumed shell subregion^12,13^ (most ventral contacts 1-2) when compared to neutral cues (Fig 1c). This power elevation was positively correlated with the patient’s ratings of drug cravings on a visual analog scale (VAS; range 1-9). The elevated low-frequency power was not observed in any other electrode contact pair along the DBS lead (Fig 1d and S1d-e), and conversely, neutral cues induced a slight suppression of power, suggesting functional and anatomical specificity consistent with this area’s subregions dissociated physiology^14^. Finally, this 1-7Hz frequency power elevation was not elicited by non-drug incentive cues during the MID and UFAT tasks (Fig S2a-b and Fig S3). Thus, low-frequency (1-7Hz) power in the NAc was hypothesized to serve as a biomarker for drug cue-reactivity in SUD.

Next, a systematic stimulation assessment was conducted across the DBS electrode contact pairs akin to standard clinical practice, but with additional focus on drug-related cravings in this subject (Fig 2a). The optimal stimulation site with the widest therapeutic window aligned with selection of the presumed NAc shell as the target, with reduction of cravings noted at the lowest dose required without adverse effects. Positive effects were also noted in more dorsal contacts in the vALIC (contacts 4+/5-) where he was previously stimulated, although higher stimulation current (8mA) was required to elicit positive effects during the evaluation. Sham-controlled stimulation suggested the acute effects of stimulation observed were not placebo effects (Supplemental Video).

**Figure 2.**
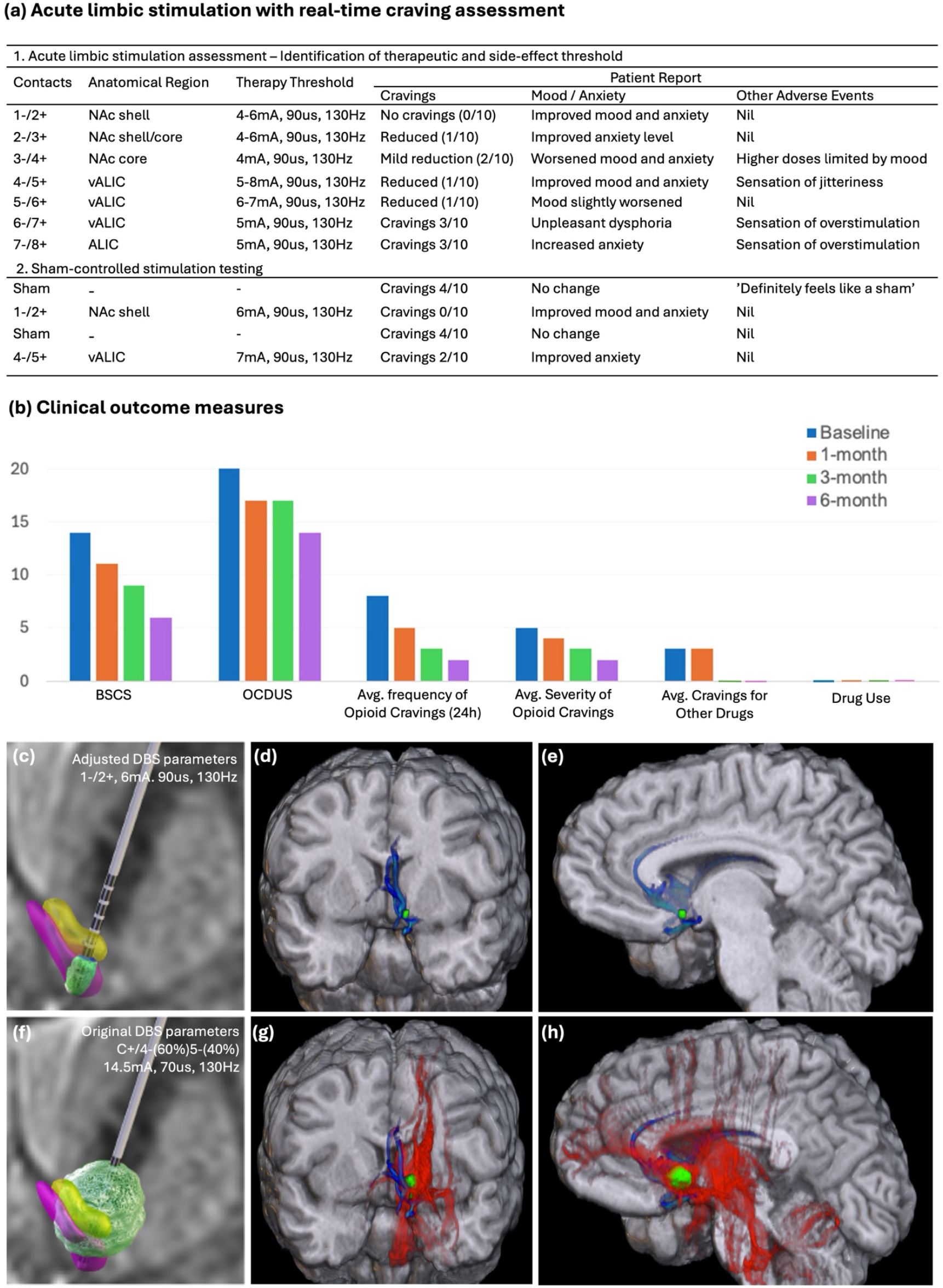
(a) Acute limbic stimulation assessment corroborated optimal stimulation settings with electrophysiological findings in the NAc shell. (b) Clinical outcome measures after continuous stimulation at 6mA reduced average craving frequency and intensity over six months. The subject remained abstinent from drug use throughout follow-up period. (c-h) Volume of tissue activation (VTA) modelling in green based on finite-element modeling (FEM) and structural connectivity analysis based on (c-e) new DBS parameters (1-/2+, 6mA, 90*μ*s, 130Hz) and (f-h) original DBS parameters (C+/4-(60%)5-(40%), 14.5mA, 70*μ*s, 130Hz). Middle and right panels display coronal and sagittal views of structural connectivity of estimated VTAs to other brain regions. Blue streamlines are connectivity associated with NAc shell stimulation. Red streamlines are connectivity associated with ALIC stimulation. *BSCS = Brief Substance Cravings Scale; DBS = deep brain stimulation; NAc = nucleus accumbens; OCDUS = Obsessive-Compulsive Drug Use Scale; VTA = volume of tissue activation*.

We then investigated the effect of NAc shell bipolar stimulation during the drug cue-reactivity task. As stimulation artifacts precluded monitoring iEEG signals simultaneous with stimulation, DBS pretreatment was delivered for ten minutes (1-/2+, 6mA, 90µs, 130Hz) prior to administration of the task, and an additional 15s stimulation period was delivered before each cue presentation (Fig 1b). This interdigitated approach allowed for the observation of stimulation effects without introducing simultaneous artifacts. Acute NAc shell stimulation attenuated the drug-cue evoked elevation of 1-7Hz low-frequency band power towards a power suppression as seen in neutral cues (\**p=*0.039, Student’s t-test), and this also correlated with >50% reduction in mean subjective rating of opioid-related cravings (6.93 vs 3.09, *p*=4.32e-7, Student’s t-test) (Fig 1e-f).

Based on these clinical and electrophysiological findings, we personalized the chronic DBS parameters (1-/2+, 6mA, 90µs, 130Hz) that required less than half of the original stimulation dose (14.5mA). Six months following this personalization, the patient reported sustained reduction in drug cravings and continued abstinence from opioid use, meeting DSM-V criteria for early remission (Fig 2b). He also reported improvement in inhibitory control and greater ease and ability to resist unwanted cravings for substances. These clinical improvements were specific to drug-related behavior, and did not affect his self-reported anxiety, mood and energy levels (Fig S4). Structural connectivity analysis of the two stimulation sites revealed that the original stimulation setting had a much larger (25x) volume of tissue activation (VTA) (896mm^3^ vs 35.4mm^3^), overlapping with the presumed NAc shell subregion, with additional, non-specific activation of other brain regions. Conversely, more apparently focal DBS delivery to the NAc shell appeared to engage a known circuit involving the subcallosal gyrus^15,12,13^ (Fig 2c-h). A basic modelling of charge requirement with this new stimulation paradigm predicted substantially improved battery lifespan (Energy Use Index 12.3 vs 39.3) of the patient’s IPG, further emphasizing the value of precision targeting and personalization of DBS therapy. These results suggest that our personalized strategy delivered DBS with similar therapeutic efficacy, fewer off-target activations, and less device charge drainage.

Here, we leveraged a rare clinical opportunity to optimize DBS therapy by understanding the electrophysiological underpinnings of opioid cravings. Patient-specific drug cue-reactive low frequency (1-7Hz) oscillatory activity localized to the presumed human NAc shell personalized a sustainable DBS intervention. This novel, serendipitous case establishes a foundation for developing individualized DBS approaches for SUD. There have been very few reports of iEEG recordings in individuals with SUD. In an intraoperative setting, prominent resting-state theta (4-8Hz) and alpha (8-14Hz) frequency activity of the ALIC and NAc were identified in patients with heroin addiction^16^. This was, however, not correlated to moment-to-moment craving states. Valencia-Alfonso et al reported an identification of a gamma-band (40-60Hz) response in the right dorsal ALIC that was associated with drug-related stimuli in a stimulation-naïve patient with heroin addiction, although this was not used to guide stimulation^17^. We present a novel paradigm of pairing a personalized cue-reactivity task with NAc region iEEG recording, which enabled identification of an anatomically and drug-cue specific electrophysiological marker that was positively correlated with craving intensity. This electrophysiologic hypersensitivity to drug-related cues correlated well with the subject’s report of increased cravings despite nearly four years of behavioral abstinence from drug use. This discovery guided a redirection of DBS programming towards the presumed NAc shell subregion, well-known to be involved in neural adaptations underlying drug-related behavioral sensitization^18^. This resulted in sustained drug abstinence and restoration of effective suppression of drug cravings at a lower charge requirement. Similar electrophysiological approaches have been employed to successfully identify biomarkers of compulsion in subjects with obsessive-compulsive disorder and loss-of-control eating^19–22^. Notably, there appears to be shared low frequency, electrographic representation of compulsion within the NAc transdiagnostically that appears to be conserved across species^23^.

Based on results of personalized tractographic structural connectivity analysis, it is postulated that the positive effects on opioid cravings from the original DBS stimulation parameters may be due to effects of engaging the ventrally-located NAc network. The presence of a drug-cue reactive electrophysiological response in the NAc is not unexpected, but the specificity of the identified biomarker presents an exciting therapeutic concept. The NAc is a central node in the mesocorticostriatal circuitry and is involved in all neurobiological stages of addiction^24^. Effects of disrupting the NAc activity with DBS have been demonstrated in rodent models of relevant endophenotypes of SUD^25–28^ and subsequently translated to off-label clinical application in humans. A recent systematic review reported that the majority of 26 human studies (71 subjects) targeted the NAc for the treatment of various SUDs^7^. While efforts have been made to improve targeting with advanced imaging techniques^29,30^, none of the studies routinely utilized a biomarker-driven approach or cue-reactivity task to optimize DBS therapy. DBS for SUD remains investigational and reported outcomes are highly variable, emphasizing the need for objective data to both guide programming and to improve understanding of addiction neurobiology^8^. This is the first demonstrated use of electrophysiological target engagement in a patient with SUD to optimize DBS therapy. Our case study also adds significant support to the potential feasibility of personalizing future SUD DBS therapy using craving-related electrographic biomarkers recorded in the NAc region to improve outcomes.

Despite the constraints of this clinically driven scenario, we observed a strong relationship between the cue-reactive 1-7 Hz low-frequency band power detected in the NAc and self-reported cravings. In the past decade, interest in brain signal-based closed-loop neurostimulation has gained significant traction in movement disorders^31,32^, epilepsy^33^, and neuropsychiatric disorders^21,34,35^. This mode of stimulation is particularly intriguing for conditions that exhibit dynamic symptomatology commonly considered for epilepsy^36,37^, but similarly, SUD^38^. Chronic and continuous DBS may in fact result in tolerance or undesirable consequences such as blunting of normal physiological functions that utilize the same pathway^39,40^. While the intent of this case study was not to facilitate a responsive stimulation strategy (in part due to limitations of the *in-situ* DBS system), the successful identification of a behaviorally relevant electrophysiological biomarker that can be modulated with acute stimulation heralds promise in utilizing electrophysiological localization of aberrant activity to guide DBS for SUD. Finally, the overall charge density delivered was substantially reduced, further providing support that this strategy may help reduce adverse and untoward effects of DBS and provide a more durable solution.

We acknowledge the limitations of this N-of-1 case study, for which the results may not be broadly generalizable to all SUD populations. In addition, iEEG recordings were constrained by the existing DBS electrode contacts. Based our findings, the low-frequency band power was strongest in the ventral-most contacts located in the NAc shell subregion. It may be possible that this electrophysiological signal be in other related brain regions not sampled by the in-situ electrode. However, we outline a compelling utility of electrophysiological guidance to optimize DBS programming and better personalize this therapeutic. We propose this cue-reactive electrophysiological signal in the NAc region as a potential candidate biomarker that warrants further investigation in ongoing and future studies of DBS in SUD.

## Online Methods

### 1. Clinical History

At the time of the current presentation, the patient was a 25-year-old male with DBS implanted 4.5 years prior. He first developed an addiction to opioids from thirteen years old following rapid withdrawal of prescription opioid medication. He also had co-morbid depression, anxiety and attention deficit hyperactivity disorder, and was treated with multiple anti-depressants, mood stabilizers and anti-psychotic medications. He started intravenous use of opiates at age fifteen and experienced episodes of opioid overdoses. He also developed disabling tardive dyskinesia that disrupted all his activities of daily living and led to severe disability (Global Assessment of Functioning Score 40) and dropped out of school. In spite of chronic methadone maintenance therapy (maximum dose 320mg/day), he had recurrent opioid cravings and required several inpatient hospitalizations for detoxification. After several years of medication optimization without success for his neuropsychiatric conditions, he was finally referred by his movement disorder specialist and psychiatrist for off-label bilateral DBS of the globus pallidus interna (GPI) to treat his severe tardive dyskinesia^41^. At the same time, DBS therapy for his severe, medication-refractory OUD was discussed. After discussion with the patient, with his parents and after careful consideration of potential risks and benefits, a consensus decision was made for bilateral GPi DBS (with right-sided IPG) to treat his tardive dyskinesia, as well as a unilateral left NAc DBS (left-sided IPG) targeted at treating his OUD (Fig S1). A unilateral NAc implant was chosen to mitigate the risk of this off-label surgery, recognizing an additional lead could be placed in the future if deemed necessary.

The DBS therapy was programmed based on standard clinical monopolar assessment used in movement disorders and psychiatric practice. Bilateral GPI DBS improved his dyskinetic movements significantly, which remained stable and managed at an outside facility with minimal adjustment required since initiation. The left NAc DBS electrode was activated based on in-clinic assessments of mood, anxiety and energy, but required a very high dose of stimulation (14.5mA) between contacts 4 and 5, which were located in the ventral anterior limb of the internal capsule (vALIC) (Fig S1c). When stimulated at yet higher doses, he experienced diaphoresis and myoclonus. Unfortunately, he was unable to tolerate a rechargeable system, which destabilized his clinical condition whenever the IPG was inadequately charged, leading to the team to recommend switching to non-rechargeable IPGs. The high dose requirement, however, led to rapid charge depletion of the IPG. Several months after clinical stabilization, he relocated to France, where a local psychiatrist managed his condition and DBS programming. In the interim, he developed episodes of hospital-acquired infections, complicated by infective endocarditis that required five cardiac surgeries for valve replacements. He abstained from drug use for several years (sustained remission) while the DBS was on, and craving levels were minimal. He was able to regain function in life, completing high school education and was able to work.

A subsequent wound erosion over the left subclavicular incision warranted removal of the left DBS IPG (extension wires were retained) and wound debridement, which was followed by a rapid recurrence of opioid cravings and drug-related thoughts (Fig 1a). The subject then re-presented to our team for further management. Due to clinical suspicion for ongoing wound infection given ecchymosis at the subclavicular site, the retained extension wire was explanted and the DBS lead externalized with an extension cable while local and blood cultures and antibiotic treatment ensued. During this period, personalized computer-based cue-reactivity tasks were performed to identify and localize a potential electrophysiological biomarker of drug cue-reactivity by recording brain activity from the DBS lead. We hypothesized that prior charge density requirements were due to non-specific engagement of the NAc region from vALIC stimulation site. To identify sites within the NAc region where cue-reactivity was most robust and specific to relevant drug cues, three computer-based tasks were conducted: a primary drug cue-reactivity task (designed with his personalized drug use habits) to assess for presence of drug-related biomarker^6^ (Fig 1b), the Monetary-Incentive-Delay, MID^10^ and the Urge-For-Action Task, UFAT^11^ (online methods, Fig S2 and S3). The two latter tasks were used as behavioral controls to assess specificity of electrophysiologic drug-cue reactivity.

There were several concerns in the management of this patient. First, a time-sensitive restoration of his DBS therapy was considered to be crucial, as he reported exacerbation in his cravings (though he had demonstrated sustained drug abstinence), a daily struggle for him. Second, the high charge requirement and intolerance of a rechargeable system was worrisome in view of his young age. Thirdly, it was critical to ensure complete eradication of the infective organism before re-implantation of the IPG due to risk to both the DBS device as well as his prosthetic heart valves. With these considerations in mind, the patient was hospitalized for treatment, with a motivation to restore his DBS therapy as soon as he was cleared by infectious disease and potentially optimize programming parameters. Due to suspicion of unresolved infection of the left IPG site, a wound exploration and swab were performed, while explanting the retained extension wire. This revealed persistence of *Staphylococcus aureus*. Upon consultation with infectious disease specialist, he was commenced on intravenous antibiotics and the DBS system was externalized percutaneously with an extension wire using previously reported surgical methods in an attempt to preserve the proximal system while antibiotic treatment ensued^42^. Using the externalized system^43,44^, we devised a plan to use personalized computer-based cue-reactivity tasks to attempt to identify potential electrophysiological biomarker of drug cue-reactivity. The overarching purpose of this was to localize the most active node encompassed by the NAc electrode for clinical purposes. We hypothesized that prior high amplitude requirements were due to non-specific stimulation target engagement and that a more precise stimulation may help reduce charge requirement and extend device lifespan which would be critical in the clinical management of this patient.

### 2. Computer-based Tasks

(a) Personalized cue-reactivity paradigm task^3,6^ The patient was interviewed by an expert in cue-reactivity (A.R.C) for specific details related to his drug use habits and preferences prior to the task. The video clips precisely reflected his paraphernalia, preferred opioid drug and injection preparation. The patient was presented with a series of 6-second-long videos in a quasi-random block design in two categories (drug cues and neutral cues). After each video presentation, the subject was asked to rate his level of craving from 1 (no craving) to 9 (extremely high craving). Eighteen repetitions of each condition were conducted to allow for comparison of responses to identify differences in electrophysiological responses to drug cues and neutral cues (control).
(b) Monetary Incentive Delay (MID) task^10^. The MID task is a validated and widely-used cognitive task to identify neural substrates of monetary reward anticipation and outcome, and has been used by this team to activate cue-reactivity within the NAc using both functional magnetic resonance imaging and electrophysiology^23,45^. The patient was presented with a visual shape cue that signals the possibility of a reward (potential gain), loss (potential loss) or neutral (no gain or loss) trial. Following the visual shape cue presentation of two seconds, a delay phase of two seconds is imposed before a target appears to prompt a response by the participant with a button press as quickly as possible to gain the reward or avoid a loss. Finally, in the feedback/outcome phase, the participant receives performance feedback with cumulative earnings.
(c) Blink Suppression Task (BST)^11^. This task uses eyeblink suppression as a model for sensory-based urges, with blocks of free blinking (30-seconds) alternating with longer blink suppression periods (60-seconds). During the free blinking periods, the instruction ‘Normal’ is displayed and subject is allowed to blink as normal. During the blink suppression period, the instruction ‘Hold’ is displayed and the participant is told to withhold blinking as long as the instruction is present. After this block, there is a recovery period with instruction ‘Ok to blink’ where the participant is allowed to blink as much as he wished, and a visual-analog-scale (VAS) rating of the urge to blink during the ‘hold’ period is collected. Eight blocks of each period are presented.

### 3. Electrophysiological Recording and Signal Analysis

After explant of the pulse generator, a Boston Scientific DBS extension wire (model NM-3138-55) was connected to the in-situ Boston Scientific DB-2201 8-ringed contacts DBS electrode, tunneled and externalized to allow intravenous antibiotic treatment of the wound infection and preservation of the remaining DBS system. An in-house customized adapter was used to connect to the Boston Scientific DBS extension wire (Boston Scientific, Marlborough, MA) to the g.HIAMP PRO system (g.tec medical engineering GmbH, Austria) for continuous intracranial electroencephalographic (iEEG) recordings of the NAc region during conduct of the computer-based tasks.

Raw iEEG data acquired at 1200Hz sampling rate was exported post-hoc for offline analysis using custom MATLAB scripts, applying a high-pass filter (1Hz) and notch filter at line noise frequency (58-62Hz) and harmonics. A time-frequency analysis using multi-taper wavelet convolution methods was used to characterize spectral power components of the cue-reactive signals. The power was extracted by squaring the magnitude of the complex Fourier-spectra and transformed by the log function. The time window length was shortened (3 to 0.13s) with increase in frequency (1-115Hz) and time step of 0.02s. The frequency resolution was dense in the low frequencies starting from 1Hz and more sparse in the high frequencies up to 115Hz using logarithmic space. The power spectrum was corrected using a common baseline from the interval 0.1-1sec before the cue was presented. Difference in mean power spectrogram obtained from each condition epoch (e.g., drug-cue or blink suppression) is compared to a neutral condition epoch (e.g., neutral-cue or normal blink) which acts as the control condition, and is then calculated for statistical significance (independent sample t-test and cluster-based permutation approach with a t-value threshold of 0.05, 1000 permutations, and an alpha level of 0.05) to identify the candidate biomarker. This was systematically performed for different frequency bands ranging from 1-115 Hz, with different size ranges.

### 4. Neuroimaging Analyses

High resolution pre-operative MRI and post-operative CT head were available for imaging analyses. T1, T2, and fast gray matter acquisition T1 inversion recovery (FGATIR) sequences were acquired at 1mm thick slices on a 3T GE scanner (GE Healthcare, Chicago, IL), as well as multi-shell 64-directions diffusion weighted imaging for tractography. The post-operative CT was co-registered to the pre-operative MRI using a 2-step linear and non-linear registration using Advanced Normalization Tools^46^. Probabilistic tractography was performed using FMRIB Software Library (FSL)’s Probtrackx2 using the NAc as a seed and the subgenual cortex as a target to subsegment the NAc into shell and core regions using clustering of streamline density according to our previously published methods^13^.

Volume-of-tissue activation (VTA) analysis was performed using Lead-DBS software based on active DBS parameters using its in-built finite element modeling (FEM)^47^. Structural connectivity analysis with streamlines was performed using the DSI-Studio^48^ using the estimated VTAs as seeds in the whole brain tractography derived from patient’s diffusion MRI.

### 5. Limbic Stimulation and Craving Assay Testing

Using the Boston Scientific external stimulator, a systematic stimulation assessment was performed across adjacent electrode contact pairs in bipolar fashion. Acute limbic stimulation effects were recorded for therapeutic as well as side-effect thresholds. The patient’s mood, anxiety and affect were recorded per standard limbic stimulation assessment. In addition, opioid craving levels were recorded using a VAS scale compared to baseline level. To confirm effects of stimulation, a short sham-controlled (patient and assessor blinded) stimulation session was also conducted to verify the effects of active stimulation.

### 6. Clinical Outcomes

The Brief Substance Craving Scale (BSCS) and Brief Addiction Monitor (BAM) questionnaires were administered to trend the levels of cravings as well as substance use over time.

## Declaration of Interests

*C*.*H*.*H has patents related to sensing and brain stimulation for the treatment of neuro-psychiatric disorders in general (USPTO serial number: 63/170,404 and 63/220,432; international publication number: WO 2022/212891 A1) as well as use of tractography for circuit-based brain stimulation (USPTO serial number: 63/210,472; international publication number: WO 2022/266000). He is a consultant for Boston Scientific, Abbott, Medtronic, and Insightec and receives honoraria for educational lectures. K*.*W*.*S is a consultant for J&J. N*.*R*.*W is a named inventor on Stanford-owned intellectual property relating to accelerated TMS pulse pattern sequences, neuroimaging-based TMS targeting, and novel psychedelic intervention for neuropsychiatric disorders; he has served on scientific advisory boards for Otsuka, NeuraWell, Magnus Medical and Nooma as a paid advisor; he also has equity/stock options in Magnus Medical, NeuraWell and Nooma. R*.*M*.*R. is a consultant for NeuroPace and receives honoraria for educational lectures as well as research grant support from Medtronic. A*.*R*.*C, C*.*H*.*H*., *D*.*W*.*O, K*.*W*.*S, L*.*Q*.,*Y*.*N. are inventors on a provisional patent application (Application No. PCT/US2024/052096) focused in SUD and related electrophysiology. The other authors declare no competing interests*.

## Acknowledgments

C.H.H is supported by the National Institute of Health (5UH3NS103446-02 and U01NS117838) and the Foundation for OCD Research. N.R.W is primarily supported by grants from the NIH (R01MH122754, UG3NS115637, R01MH128311, R01MH118388), the Pritzker Neuropsychiatric Disorders Research Consortium (24490), the New Venture Fund (010038-2020-06-01, 011665-2020-08-01), the Wellcome Trust (220839-1), and funds from Bayshore Global Management. The authors would like to thank Dr Robert Melanka for his critical review of the manuscript, and above all, the patient for his participation and commitment to this novel, clinical application.

## Notes

### Summary of Updates

Updated figures to accurately reflect results

