## Supplemental Data for "Localizing electrophysiologic cue-reactivity within the nucleus accumbens guides deep brain stimulation for opioid use disorder"

### Supplementary Figures / Extended Data Figures

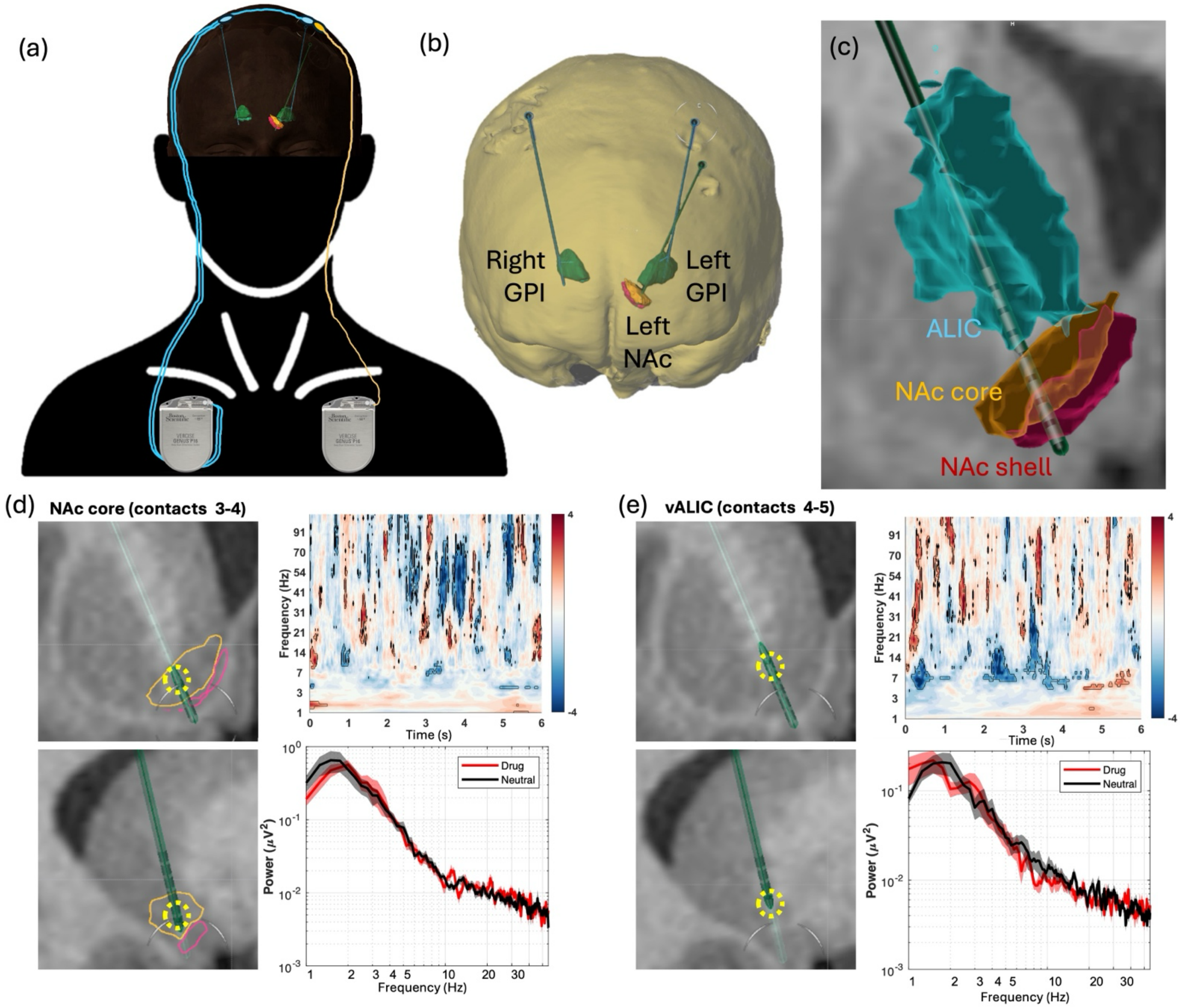


**Figure S1.** (a) The patient received bilateral GPI DBS to treat generalized tardive dyskinesia, as well as left NAc DBS to treat opioid use disorder (OUD). The bilateral GPI DBS were connected to a right-sided subclavicular Boston Scientific Vercise PC DB-1140 pulse generator (IPG), denoted in blue while the left NAc DBS electrode was connected to a separate left-sided subclavicular Boston Scientific Vercise PC IPG (yellow). (b) An expanded 3-dimensional view of the intracranial DBS electrode positions in the subject. The Boston Scientific directional DB-2202 electrodes were implanted in bilateral GPI targets, while the eight ring-contacts Boston Scientific DB-2201 electrode was implanted in the left NAc. (c) Expanded coronal view (*surgical view*, i.e. left is on the left) of the left NAc DBS electrode showing locations of each DBS electrode contact. 3-dimensional segmentations of the NAc shell (pink), NAc core (yellow) and anterior limb of the internal capsule, ALIC (blue) are shown as segmentation overlays. (d) Left; Coronal (top) and Sagittal (bottom) MRI views of the location of electrode contacts 3-4 in the NAc core. Right; Time-frequency analysis (top) and power spectral density plot (bottom) of iEEG recordings obtained from contacts 3-4 did not identify any significant difference in power between drug cues and neutral cue trials. (e) Left; Coronal (top) and Sagittal (bottom) MRI views of the location of electrode contacts 4-5, where his previous stimulation was set. Right; Time-frequency analysis (top) and power spectral density plot (bottom) of iEEG recordings obtained from contacts 4-5 did not identify any significant difference in power between drug cues and neutral cue trials. *ALIC=anterior limb of the internal capsule; DBS=deep brain stimulation; GPI=globus pallidum interna; IPG=implantable pulse generator; NAc=nucleus accumbens; OUD=opioid use disorder; PC=primary cell.*

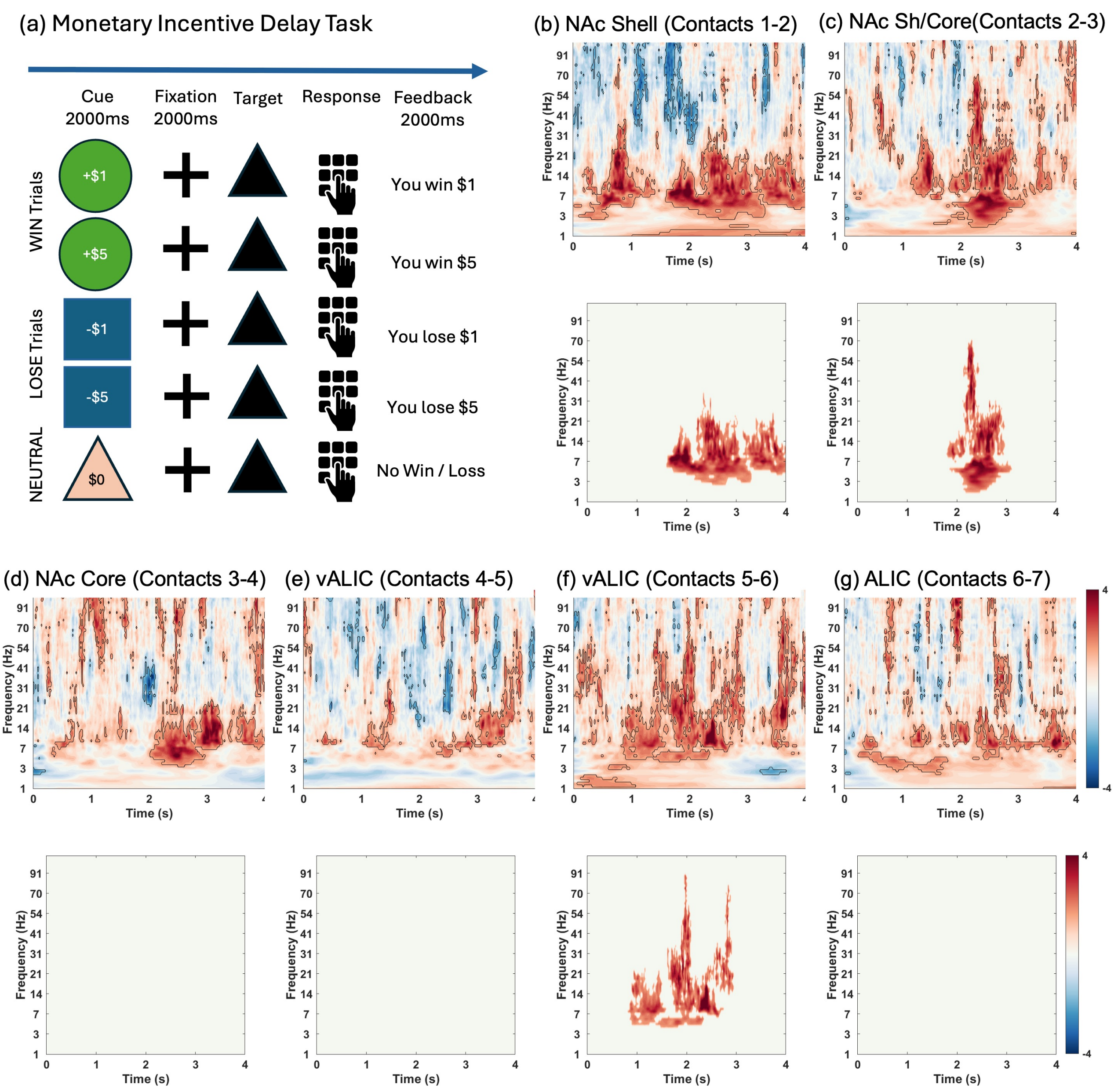


**Figure S2.** Time-frequency analysis of the Monetary-Incentive-Delay (MID) Task. (a) Schematic of the MID Task. In this task, the participant is presented with a visual shape cue that signals the possibility of a reward (potential gain), loss (potential loss) or neutral (no gain or loss) trial. Following the cue, a delay phase of two seconds (fixation) is present before a target appears to prompt a response by the participant with a button press as quickly as possible to gain the reward or avoid a loss. Finally, in the feedback/outcome phase, the participant receives performance feedback with cumulative earnings. (b-g) Time-frequency analysis of the electrophysiological recordings showed increase in power in multiple electrode contact pairs in a wider frequency range, very different from the drug-cue reactivity paradigm. Each top panel represent power spectrum difference when the patient was presented with high reward (+$5) cue compared to no win/loss ($0) cue in each contact pair as indicated. Bottom panel displays cluster-corrected differences that were significant ($p<0.01$).

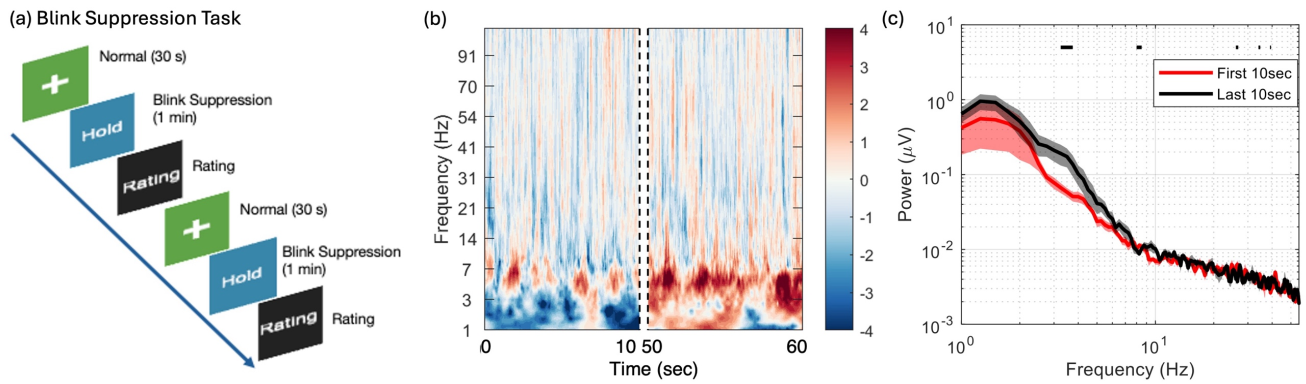


**Figure S3**. Time-frequency analysis of the Blink Suppression Task (BST). (a) Schematic of the BST. In this task, the participant is presented with blocks of free blinking (30 seconds), alternating with blink suppression blocks (1 minute) when the instruction 'Hold' is displayed and participant is told to withhold blinking as long as the instruction is present. After this block, a visual-analog-scale (VAS) rating of the urge to blink during the 'hold' period is collected. Eight blocks of each period are presented. (b) Time-frequency plot of the first 10-seconds (assumption: low urge to blink) and the last 10-seconds (assumption: high urge to blink) of the blink suppression block was compared, and did not show a significant change in power spectrum across all electrodes ($p=0.1480$). This figure shows a representative spectrogram in the NAc shell (electrode contacts 1-2). (c) Power spectrum density plot comparing the first 10 seconds (red) and last 10 seconds (black) of the blink suppression block showing increased power in a very narrow frequency band denoted with black line.

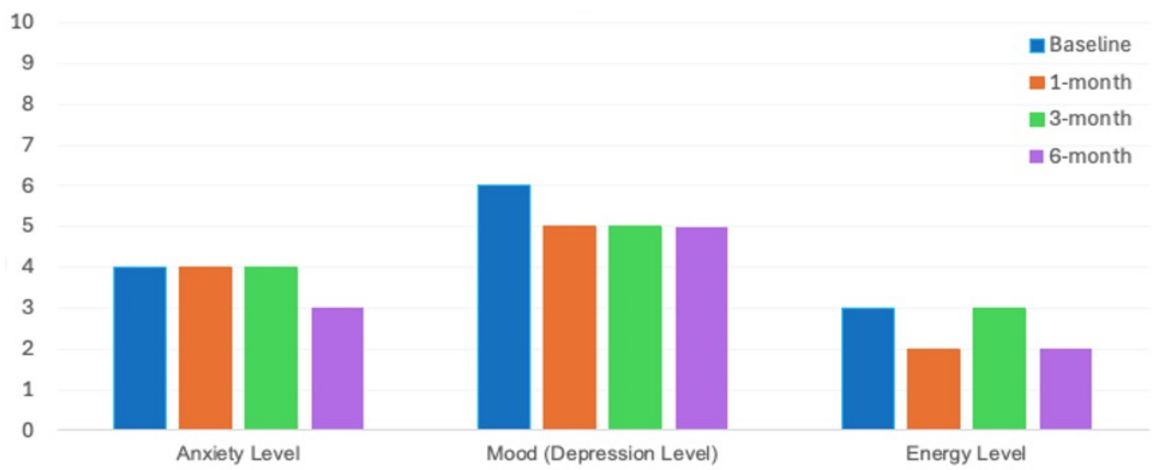


**Figure S4**. Subject self-reported ratings of anxiety, mood and energy levels over 6 months follow-up
